# Improving the learning curve in monoportal endoscopic lumbar surgery: development and validation of a porcine training model

**DOI:** 10.1101/2025.09.18.677106

**Authors:** Laura Olías-Ortiz, Rafael Llombart-Blanco, Gloria Abizanda-Sarasa, Isabel Lacave-Mas, Carmen Valverde-Gestoso, Matías Alfonso-Olmos

## Abstract

**Study design:** Experimental animal study.

**Purpose:** Endoscopic spine surgery (ESS) has gained substantial traction globally. Proficiency in ESS demands a steep learning curve and rigorous technical training. This study aims to develop and validate a porcine model as a high-fidelity, reproducible platform for hands-on training in lumbar endoscopic procedures.

**Methods:** Thirteen skeletally mature Landrace x Large white pigs underwent lumbar endoscopic decompression via a transforaminal approach at L4-L5 and an interlaminar approach at L5-L6. Specific procedural modifications were introduced to optimize anatomical correlation and surgical ergonomics. Following a seven-day postoperative period, necropsies were performed to evaluate procedural outcomes and adverse events. Operative times, intraoperative events, and postoperative outcomes were systematically recorded.

**Results:** Endoscopic decompression was successfully completed in all specimens. Mean operative time was 134.7 ± 32.3 minutes. Intraoperative complications occurred in 4 animals (30.8%): uncontrollable epidural bleeding (case 1), dural tear (case 7), technically difficult transforaminal access in a low-weight pig (33.8 kg, case 8), and a complex interlaminar approach leading to spinal cord injury (case 13). One additional animal (case 4) developed paraplegia without structural findings at necropsy, attributed to intraoperative positioning. Necropsy revealed asymptomatic epidural hematomas in three cases (9, 11, 12). Three animals (23.1%) required early euthanasia based on humane criteria. A significant learning curve effect was demonstrated, with operative time decreasing over the series (ρ = –0.73, p = 0.004).

**Conclusions:** The porcine lumbar spine provides a reliable and anatomically valid model for ESS training. Awareness of species-specific anatomy, specimen size, and positioning is essential to maximize reproducibility and safety.

## INTRODUCTION

Endoscopic spine surgery (ESS) has emerged as a minimally invasive alternative to traditional open surgery, offering several advantages that include reduced soft tissue damage, lower hospitalization rates, and faster patient recovery [1, 2]. ESS facilitates earlier postoperative mobilization and a quicker return to work. Moreover, its reoperation rate is comparable to other minimally invasive techniques, such as microdiscectomy. ESS is also a valid option for elderly patients with surgical risks who suffer from spinal stenosis. However, ESS is not without risks, with overall complication rates reported as 9.26% [3], reaching up to 21% in some series. The leading countries in ESS, based on publication volume, are China, South Korea, and the United States, yet the technique is experiencing significant growth in Europe [4]. Despite increasing recognition of ESS and its benefits, surgeons face adoption challenges, mainly due to limited funding for specialized training, a lack of hands-on courses, and the high cost of equipment, factors that collectively increase procedural expenses [5]. These training limitations represent a critical obstacle to the safe implementation of ESS. The factors mentioned above substantially contribute to the difficulty of overcoming the ESS learning curve, necessitating innovative training approaches to ensure safety and competence. The technically demanding nature of ESS, combined with its limited visualization and confined working space, requires structured, progressive training before clinical application. In this context, developing effective training models becomes fundamental. Given the ethical, logistical, and financial limitations associated with cadaveric models, large animal models have become essential tools for the development and training of ESS techniques. Model selection is typically based on anatomical similarity to the human spine and surgical feasibility.

The choice of the porcine model is supported by multiple anatomical and practical advantages. Among the available options, the porcine model is the most commonly used, particularly due to its comparable vertebral architecture and intervertebral foramen morphology [6]. Anatomical studies have shown that the porcine lumbar spine exhibits vertebral dimensions, intervertebral-disc height, and spinal-canal structure that closely approximate human characteristics, providing a realistic training environment for endoscopic procedures.

Moreover, the porcine model offers practical advantages over other animal alternatives. Ovine and bovine models are also employed for selected spinal or cranial procedures, though with more restricted applicability [7]. The porcine model permits practice of both main endoscopic approaches—transforaminal and interlaminar—offering versatility in surgical training. Its availability, cost-effectiveness, and favorable ethical considerations make it the most practical option for structured training programs.

Despite known anatomical differences, the porcine model retains its validity for training. Although anatomical differences—such as reduced disc height and narrower interlaminar windows—have been reported in the porcine spine [8,9], these variations can be accommodated through technique adaptation, rendering the model highly reproducible and cost-effective. Previous anatomical studies have documented these discrepancies [10], yet there remains a need to standardize the porcine model specifically for monoportal endoscopic decompression and to assess its practical suitability for procedural training.

Therefore, the aim of this study was to validate the porcine lumbar spine as an experimental model for monoportal endoscopic decompression, assessing its anatomical fidelity, technical feasibility, and applicability for structured surgical training programs.

## MATERIAL AND METHODS

### 1. Animal and housing conditions

Thirteen skeletally mature Landrace × Large White pigs (6 males, 7 females) were enrolled. Selection criteria included a target body weight of 40–45 kg, skeletal maturity, and absence of locomotor or neurological deficits on preoperative veterinary examination. This weight range was chosen because it provides vertebral dimensions and anatomical landmarks comparable to those in the human lumbar spine. Animals of both sexes were included, as sex was not deemed to affect surgical access, anatomical correlation, or procedural outcomes in this experimental model. The animals were housed in social groups during the acclimatization period until their inclusion in the experimental process, at which time they were individualized until the end of the study period, thus allowing a correct evaluation of the animal’s condition, control of solid/liquid intake and intestinal mobility. Environmental enrichment was provided alternately (fruit ice cream, balls, porky play, string, marshmallow) and standard environmental conditions were maintained according to current regulations. All husbandry complied with European and institutional animal-care regulations, and the study protocol was approved by the University of Navarra Animal Research Ethics Committee (Project 49/23, 25 Aug 2023).

### 2. Anaesthetic protocol and peri-operative care

Food was withheld for 6–8 h pre-operatively. Premedication was delivered intramuscularly: atropine 0.01 mg kg^−1^, tiletamine–zolazepam 4 mg kg^−1^ and medetomidine 0.05 mg kg^−1^. Following sedation, propofol 1–2 mg kg^−1^ IV was administered for endotracheal intubation. General anaesthesia was maintained with 2 – 3% inhaled sevoflurane supplemented by a fentanyl continuous-rate infusion (0.03 – 0.10 mg kg^−1^ h^−1^). A single prophylactic dose of cefazolin 1 g IV was given before incision. At wound closure, methylprednisolone 1 mg kg^−1^ IV was administered to limit local inflammation. Post-operative analgesia consisted of a transdermal fentanyl patch (75 μg h^−1^) and meloxicam 0.4 mg kg^−1^ SC once daily for 3 days. Surgical wounds were cleaned daily with 2% aqueous chlorhexidine and hypochlorous solution.

### 3. Surgical Technique

The animal was placed in a prone position on a radiolucent table with its legs extended forward to promote kyphosis of the lumbar spine, thereby facilitating surgical access. The dorsal aspect of the pig was shaved, cleaned with antiseptic soap and the surgical site was prepared using an iodinated alcohol solution. Sterile surgical drapes were applied, exposing the lumbar midline. The surgery was performed under fluoroscopic guidance. All procedures were performed by a single experienced spine surgeon to minimize variability, using the same endoscopic equipment employed in human clinical practice.

#### 3.1 Transforaminal Approach (L4-L5)

A needle is first placed on the skin to identify and mark the spinous processes. The intervertebral disc L4–L5 is then located and marked on the skin using fluoroscopic guidance. In the lateral fluoroscopic view, the needle should be positioned over both the spinous process and the facet line to ensure accurate trajectory. The standard entry point is 10–12 cm lateral to the midline, depending on the animal’s size.

Under fluoroscopic control, the needle is advanced toward the target disc space, ensuring the tip is located at the facet joint in the lateral view and between the medial and lateral pedicular lines in the anteroposterior view. The target point for needle insertion was the same as in human procedures: the medial pedicular line in the AP view and the posterior vertebral or posterior disc line in the lateral view. An 18G needle was inserted into the intervertebral disc, and a mixture of 1 mL of methylene blue and radiographic contrast agent was injected into the disc space to facilitate both direct and radiological visualization (Figs 1 and 2).

**Fig 1.**
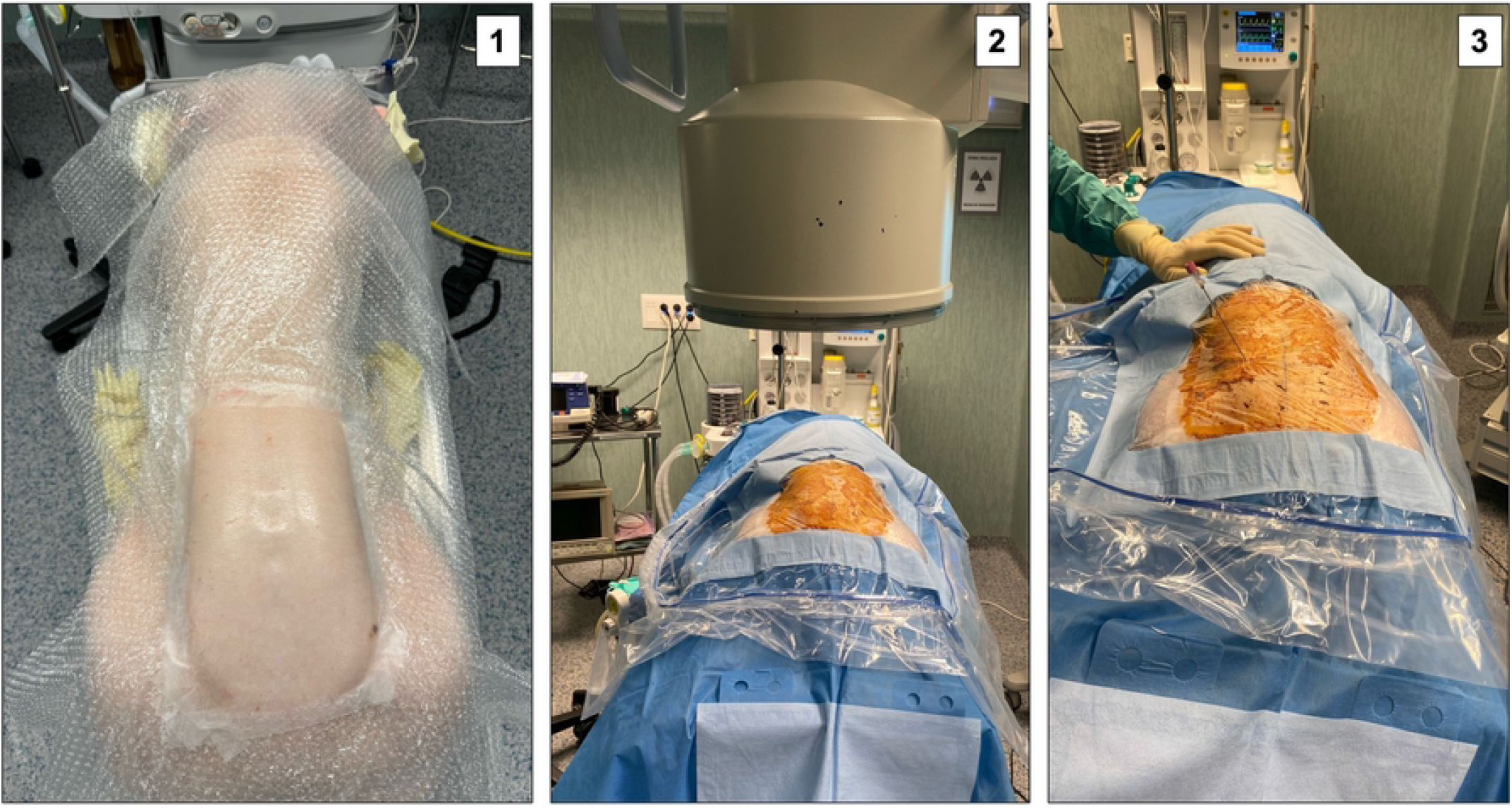
First steps of surgery. 1: Prone positioning of the animal with the hind limbs extended forward; 2: Placement of sterile drapes and fluoroscopic equipment; 3: Insertion of the guide needle for the transforaminal approach.

**Fig 2.**
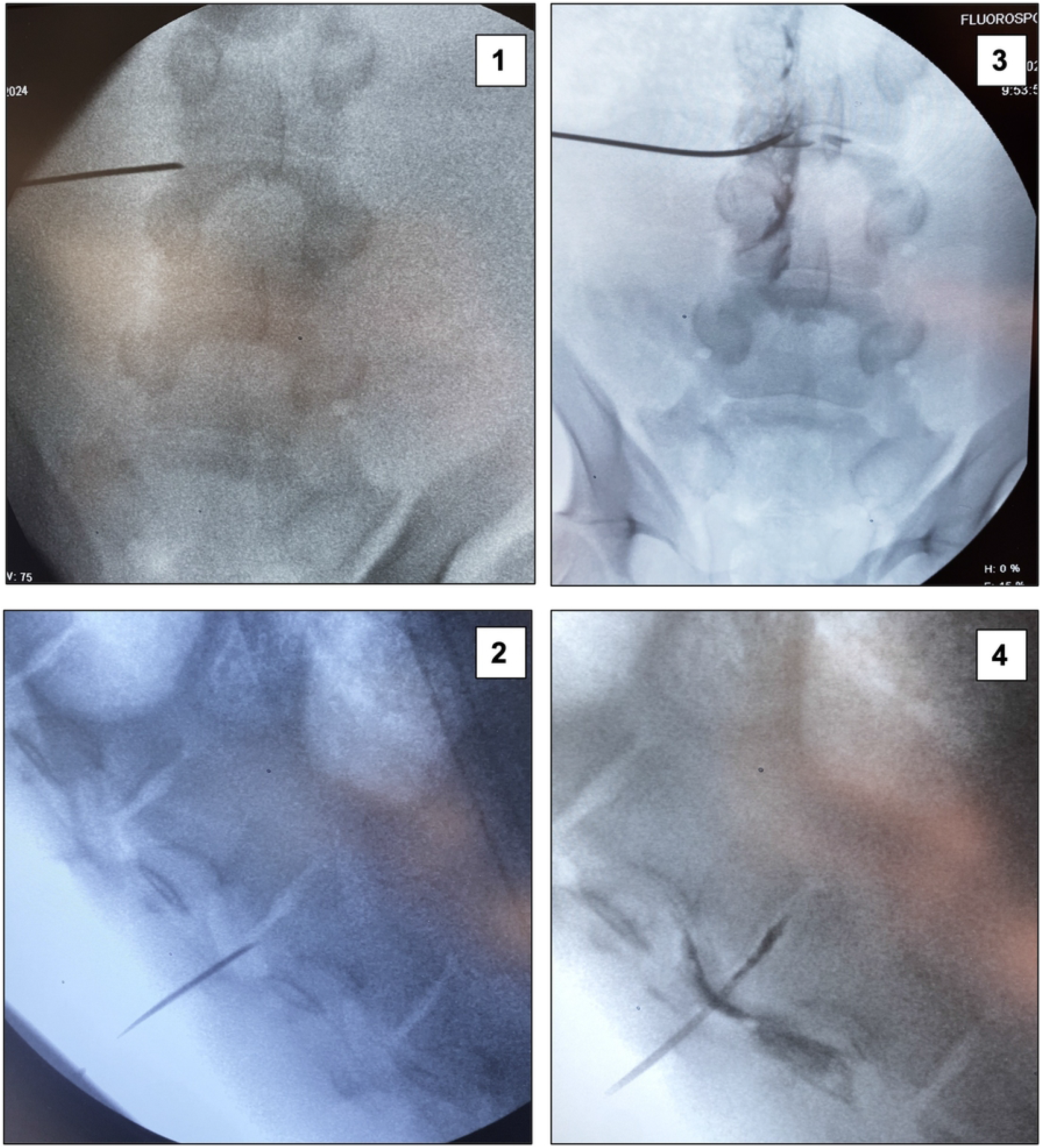
Fluoroscopic localization for lumbar endoscopic decompression. 1-2: Fluoroscopic localization of the transforaminal approach trajectory; 3-4: Discography with contrast injection into the intervertebral disc space.

A 2 cm skin incision was made at the entry site. Sequential dilators were introduced over the guidewire to establish the working channel, after which the endoscope was inserted. Instrumentation was performed using a monoportal endoscopic spine surgery system. Partial resection of the facet joint was performed using a high-speed burr to access the working channel, and bone fragments were removed with rongeurs until the foramen was adequately exposed. The procedure was considered complete when the dural sac and nerve root were clearly visualized and confirmed to be decompressed (Fig 3).

**Fig 3.**
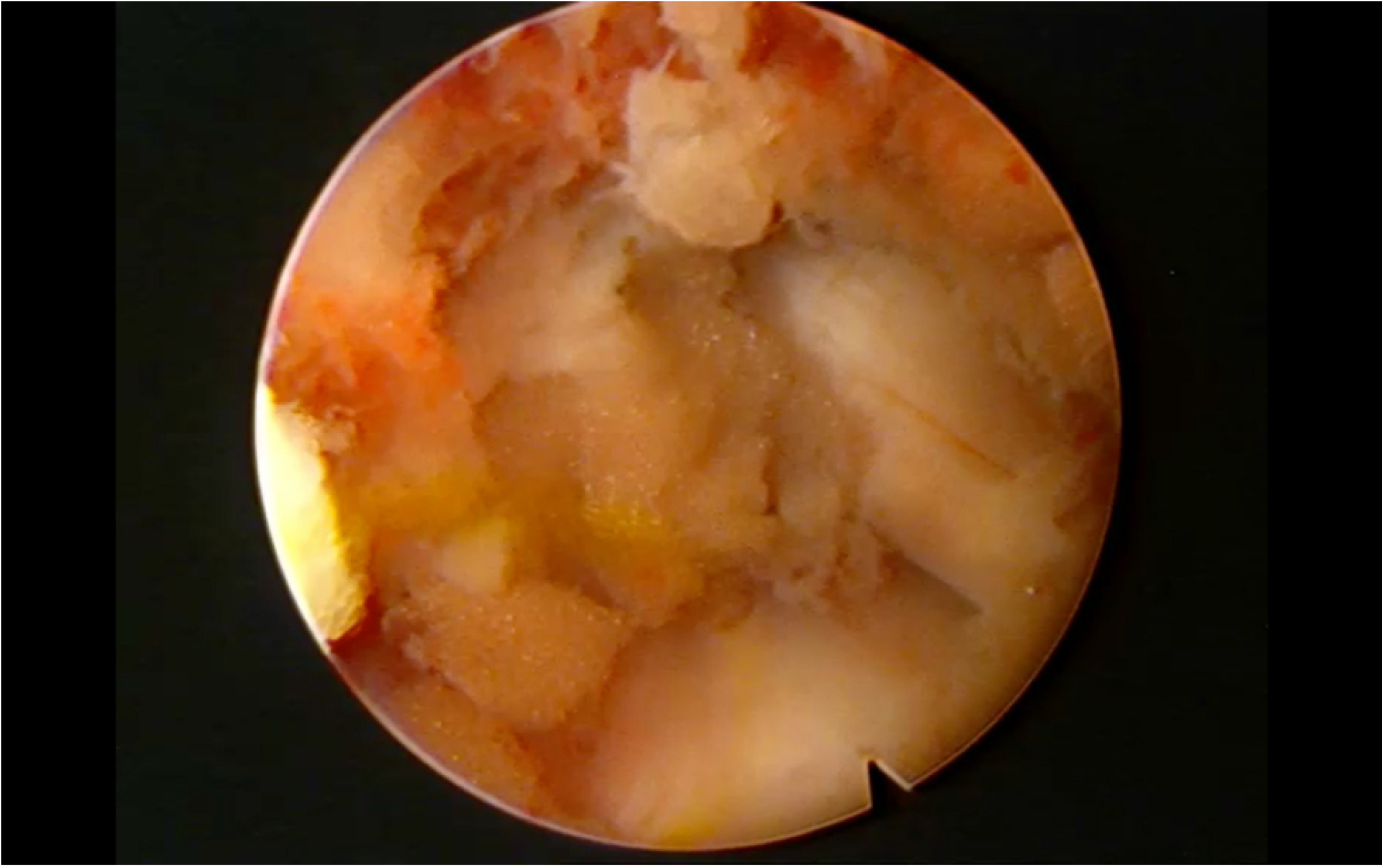
Intraoperative view of the transforaminal approach. Endoscopic visualization of the working channel after partial facetectomy, exposing the exiting nerve root and dural sac.

#### 3.2 Interlaminar Approach (L5-L6)

Under fluoroscopic guidance, the L5-L6 interdisc space and interlaminar window was identified. Entry point was as medial as possible, just lateral to the spinous process centered on interlaminar window. The endoscope was introduced, and a laminectomy was performed using a laminectomy burr. Due to small interlaminar windows, flavectomy was then performed using microscissors until the spinal cord and the adjacent nerve root were clearly visualized and decompressed (Fig 4).

**Fig 4.**
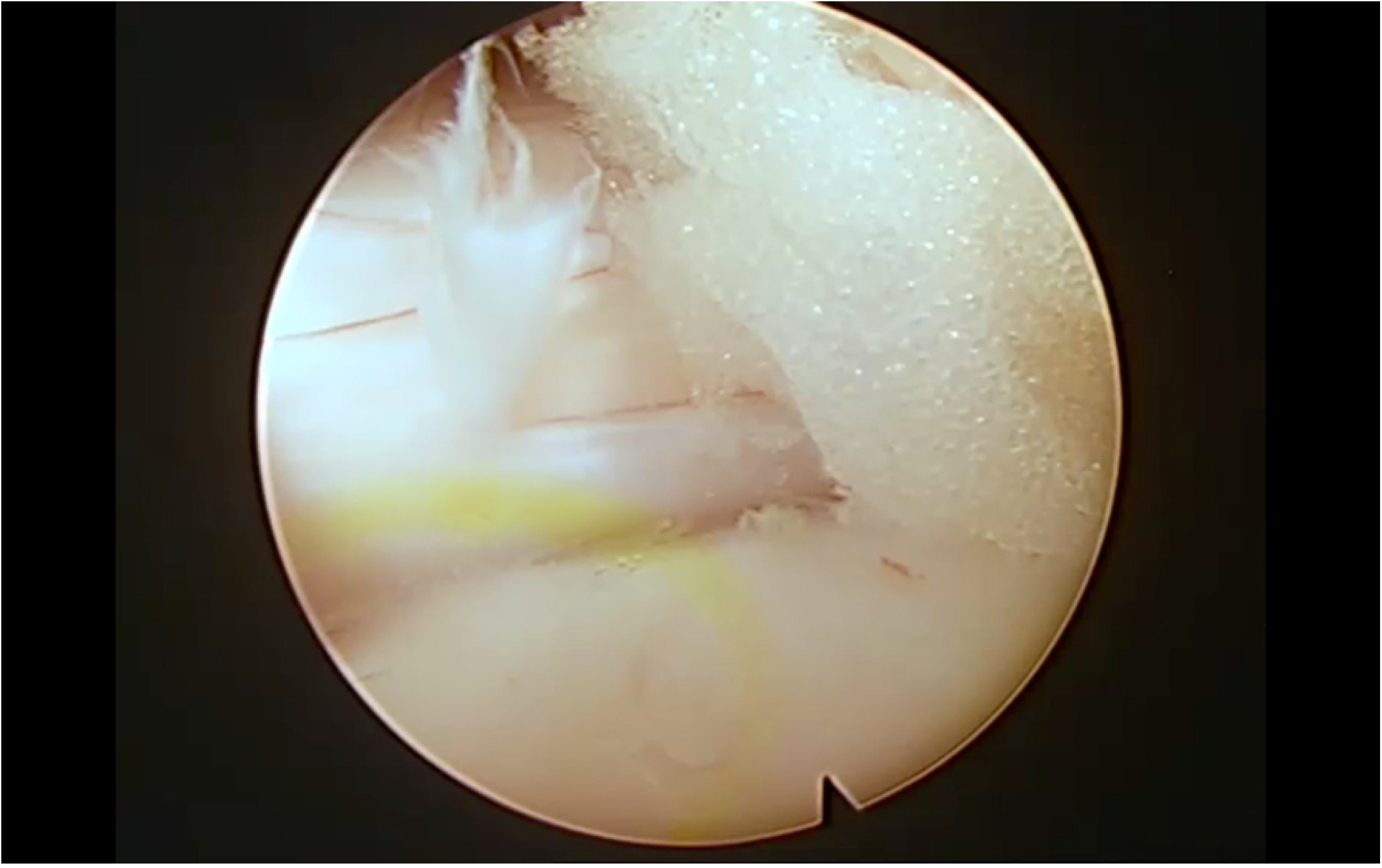
Intraoperative view of the interlaminar approach. Endoscopic decompression following flavectomy, with clear visualization of the dural sac and adjacent nerve root.

It should be noted that, compared to humans, the porcine intervertebral disc is narrower, and the nerve root emerges in a more vertical orientation.

Continuous irrigation with elevated normal saline was used to maintain a clear surgical field. Hemostasis was achieved using an electrocautery device

The skin was closed using 0 silk suture. Following wound closure, the surgical site was disinfected with an iodized alcohol solution.

### 4. Post-operative Monitoring and Functional Assessment

Animals were kept in the operating room until restoration of reflexes, spontaneous respiration and cardiovascular stability, then returned to their pens. Re-examinations were conducted at 3 hours and every 12 hours thereafter. Daily evaluations included hind-limb muscle strength, ability to stand and ambulate, pain scoring, and wound inspection. Observation frequency increased if abnormalities were detected.

### 5. Euthanasia Criteria

The primary endpoint for euthanasia was the absence of response to specific treatments for any presented symptoms. Deterioration criteria included: >20% weight loss, cyanotic mucous membranes, dyspnea, prostration, body temperature ≥40°C for >2 days, prolonged lower extremity paralysis (>3 days), absent response to painful stimuli post-surgery, or surgical wound infection with purulent exudate for >4 days. Euthanasia was elected for animals that presented with these criteria and failed to respond to therapeutic intervention

### 6. Necropsy Procedure

On postoperative day 7, animals were euthanised with intravenous KCl (2–4 mg kg^−1^) under deep anaesthesia. The lumbar spine was harvested *en bloc* to verify operative levels, assess approach accuracy and document complications such as epidural haematoma, dural tear or nerve-root injury.

### 7. Datta Collection

The primary data collected was a detailed description, with intraoperative photographs and/or video, of the surgical technique. The following parameters were also recorded:

- Surgical time (defined as the time from skin incision to skin closure)
- Intraoperative monitoring (Blood pressure, heart and respiratory rate, temperature, CO2)
- Any intraoperative complications (e.g., dural tear, nerve root injury).

### 8. Statistical analysis

Continuous variables were expressed as mean ± standard deviation (SD) or median (interquartile range), as appropriate. The relationship between case sequence and total operative time was evaluated using Spearman’s rank correlation. Statistical analyses were performed using Stata version 17.0 (StataCorp LLC, College Station, TX, USA). A *p*-value <0.05 was considered statistically significant.

## RESULTS

Thirteen pigs (6 male, 7 female) with a mean animal weight of 44.7± 5.3kg (range: 33.8–61.7 kg) underwent lumbar endoscopic decompression using the monoportal technique, utilizing two approaches: transforaminal at L4-L5 and interlaminar at L5-L6. The actual body weight at the time of surgery ranged from 33.8 to 61.7 kg, as some animals gained weight during the acclimatization period, while others arrived slightly below the target range. Surgical data, including operative times, approach-specific details, complications, and necropsy findings, are summarized in table 1.

**Table 1.** Summary of experimental surgical cases, including animal sex, total operative time, intraoperative and postoperative complications, and necropsy findings.

| ID | SEX<sup>a</sup> | WEIGHT (kg) | TOTAL SURGICAL TIME (min) | INTRAOPERATIVE COMPLICATIONS | POSTOPERATIVE COMPLICATIONS | NECROPSY FINDINGS |
| --- | --- | --- | --- | --- | --- | --- |
| 1 | F | 45.85 | 192 | Uncontrollable epidural bleeding during the transforaminal approach | Hind limb paralysis. Humane endpoint criteria met | Spinal cord injury |
| 2 | F | 47.6 | 181 | No | No | No significant findings |
| 3 | F | 43.25 | 157 | No | No | No significant findings |
| 4 | M | 52.95 | 167 | No | Hind limb paralysis. Humane endpoint criteria met | No significant findings |
| 5 | F | 40.8 | 140 | No | No | No significant findings |
| 6 | F | 36.3 | 90 | No | No | No significant findings |
| 7 | M | 39.2 | 122 | Inadvertent durotomy during the transforaminal approach | No clinical signs of motor weakness | Cerebrospinal fluid fistula |
| 8 | F | 33.8 | 110 | Technically difficult transforaminal approach. low-weight pig | Delayed ambulation | Small intramedullary hematoma |
| 9 | M | 44 | 130 | No | No | Epidural hematoma |
| 10 | M | 44.9 | 120 | No | No | No significant findings |
| 11 | M | 45.2 | 111 | No | No | Epidural hematoma |
| 12 | M | 61.7 | 116 | No | No | Epidural hematoma |
| 13 | F | 46.1 | 115 | Technically difficult transforaminal approach | Hind limb paralysis. Humane endpoint criteria met | Spinal cord injury |

<sup>a</sup> F:female; M:male

^a^ F:female; M:male

Comprehensive surgical data revealed a mean total surgical time of 134.7 ± 32.3 minutes. The transforaminal approach (TF) required a mean approach time of 53.3 ± 18.5 minutes, with a mean coagulation time of 20.1 ± 33.1 seconds and a mean saline usage of 4.8 ± 3.6 Liters. The interlaminar approach (IL) was performed with a mean approach time of 51.5 ± 17.6 minutes, a mean coagulation time of 4.1 ± 10.7 seconds, and a mean saline usage of 5.5 ± 3.6 Liters. Post-mortem examination confirmed accurate vertebral level access in all instances. Nine animals (69.2%) were able to stand on all fours after waking from anesthesia. There were no intraoperative monitoring incidents. Surgical complications were observed in 4 cases (30.8%).

- **Case 1:** Uncontrollable epidural hemorrhage during the transforaminal approach, resulting in spinal cord injury and paraplegia. The animal was euthanized on postoperative Day 4. Necropsy confirmed bilateral epidural hematomas.
- **Case 7**: Dural tear during the transforaminal approach without neurological sequelae. Necropsy confirmed an asymptomatic cerebrospinal fluid fistula.
- **Case 8:** Difficult transforaminal access in a low-weight animal (33.8 kg), associated with delayed ambulation until postoperative Day 5. Necropsy revealed a small intramedullary hematoma with mild spinal cord displacement.
- **Case 13:** Technically demanding interlaminar approach complicated by spinal cord injury, leading to paraplegia. The animal was euthanized on postoperative Day 3; necropsy confirmed focal cord contusion.

Additionally, epidural hematomas were identified in three other animals at necropsy. These findings were clinically silent, as none of the animals exhibited postoperative neurological deficits.

Only three animals (23.1%) were euthanized prior to the study endpoint according to humane endpoint criteria (Cases 1, 4, and 13). In Case 4, the animal developed paraplegia without any intraoperative complication; necropsy did not reveal structural abnormalities. This case had a prolonged time on the surgical table due to anesthetic management, and the deficit was attributed to possible positioning-related ischemia.

A Spearman’s rank correlation was performed to evaluate the relationship between surgical case number (i.e., the chronological order of procedures) and total surgical time. The analysis revealed a statistically significant negative correlation (ρ = -0.73, p = 0.004), indicating that total surgical time decreased as the number of procedures performed increased. These findings demonstrate a clear learning curve effect, with increased experience leading to improved surgical efficiency (Fig 5).

**Fig 5.**
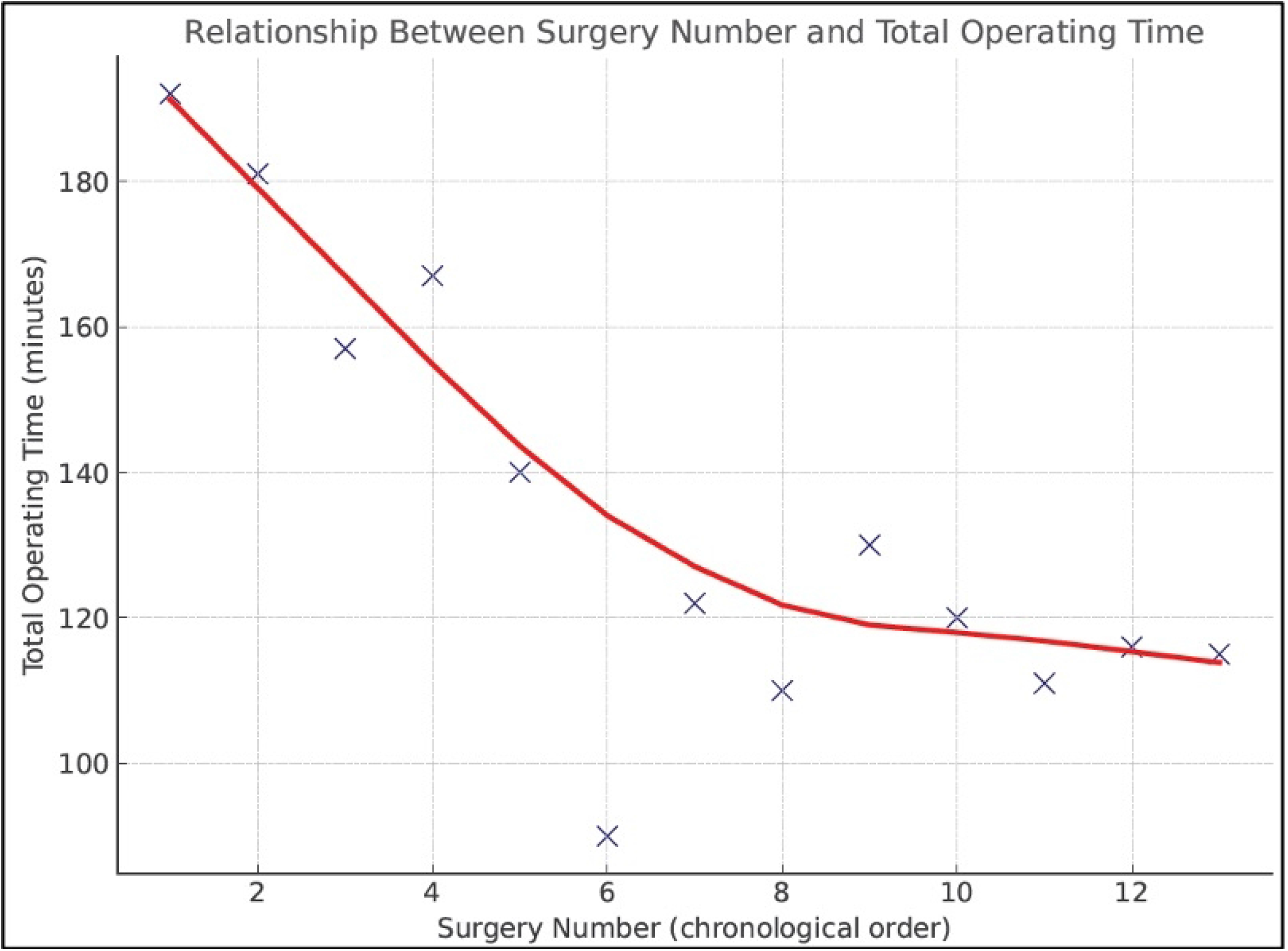
Learning curve for endoscopic lumbar spine surgery in the porcine model. Each point represents the total surgical time for a single case. The downward trend demonstrates improved efficiency with increasing surgical experience.

## DISCUSSION

This study demonstrates the feasibility and anatomical compatibility of using the porcine model for monoportal endoscopic lumbar spine surgery. In this context, the anatomical landmarks and spatial organization of the porcine lumbar spine proved sufficiently analogous to the human anatomy to support fundamental procedural steps such as transforaminal and interlaminar access. The porcine lumbar spine has been widely validated as a suitable model for endoscopic spine surgery training and research due to its anatomical similarities to the human lumbar region. Studies such as those by Amato et al. [6] and Gotfryd et al. [8] have shown that the overall vertebral architecture, neural decompression possibilities, and pedicular instrumentation techniques in pigs closely resemble those in humans. However, several anatomical peculiarities must be considered [9,10].

An additional anatomical factor relevant to surgical planning is the orientation and distribution of lumbar nerve roots. Unlike in humans, where lumbar roots follow a caudally oblique trajectory, in pigs the nerve roots emerge from the spinal cord in a more horizontal plane, nearly perpendicular to the cord. In addition, the porcine spine exhibits longer spinous processes, a thinner ligamentum flavum, and a narrower anteroposterior vertebral body diameter, which may necessitate adaptation of surgical instruments and techniques. One crucial consideration is the length of the spinal cord in pigs, which extends to the sacral region, and that they typically have six lumbar vertebrae. This anatomical characteristic increases the risk of spinal cord injury during lumbar procedures, highlighting the need for meticulous surgical technique and precise anatomical knowledge.

Surgeons training with this model must develop a precise understanding of these neuroanatomical differences to perform safe and reproducible procedures.

Our experience confirms that these anatomical differences can present technical challenges during endoscopic lumbar procedures, particularly in animals weighing less than 40 kg. In our series, the only animal below this threshold (33.8 kg) required a more complex approach due to reduced access and anatomical constraints, resulting in delayed recovery and evidence of intramedullary hematoma on necropsy. This aligns with previous literature emphasizing the importance of specimen size in ensuring adequate working space and reproducibility of surgical maneuvers. Based on our findings, we recommend the use of pigs weighing more than 40 kg for endoscopic lumbar decompression studies, as larger animals provide vertebral dimensions and tissue planes more comparable to those encountered in human surgery, reducing the risk of technical complications and facilitating the learning process.

Interestingly, in one case of paraplegia that required early euthanasia based on humane endpoint criteria (case 4), necropsy did not reveal any structural cause for the neurological deficit. The authors hypothesize that this deficit may have resulted from the intraoperative positioning, specifically prolonged hip flexion during the surgical procedure. In this case, surgical table time was extended due to prolonged anesthetic induction and stabilization. All procedures in our study were confirmed at the correct vertebral level by post-mortem examination, supporting the reliability of the porcine model for anatomical and technical validation. Nevertheless, the inherent anatomical differences—such as narrower disc spaces and lower disc heights—should be accounted for when extrapolating results to clinical practice or using the model for training purposes.

One of the most relevant findings in our study was the significant learning curve observed, reflected by a steady reduction in operative time as the number of procedures increased. This trend, supported by statistical analysis, highlights the utility of the porcine model not only for anatomical simulation but also for progressive acquisition of technical skill. The model effectively reproduces the challenges of real surgical scenarios and allows iterative refinement of technique, making it especially valuable for surgeons at early stages of endoscopic training.

Although objective assessments of motor function were systematically performed, this study is limited by its short postoperative follow-up period and the relatively small sample size, which may have restricted the detection of delayed-onset complications or long-term neurological sequelae.

Finally, while the porcine model does not fully replicate all aspects of human spinal anatomy, it remains a robust and reliable platform for developing and testing endoscopic decompression techniques. From a translational perspective, training in a porcine model bridges the gap between conceptual learning and clinical application. It provides surgeons with exposure to real biological tissues and hemodynamic responses, which are absent in most benchtop simulators. Therefore, this model not only enhances anatomical familiarity and technical confidence but also promotes the development of tactile and procedural competence that are essential for clinical translation.

Future studies should focus on establishing objective performance metrics and validating the correlation between training in this model and outcomes in human procedures. Additionally, anatomical mapping to further refine approach trajectories and reduce variability will improve standardization and educational reproducibility.

With appropriate anatomical knowledge and careful case selection, this model provides a high-fidelity, ethically justifiable alternative for surgical training and preclinical validation.

## CONCLUSION

The porcine model has proven to be a valuable and anatomically viable platform for practicing monoportal endoscopic lumbar spine surgery. Despite minor anatomical differences compared to humans, it allows for safe and reproducible execution of both transforaminal and interlaminar approaches with satisfactory visualization and spatial orientation. As such, it represents an essential intermediary between artificial simulators and clinical practice, offering biological realism without compromising safety.

## ETHICAL CONSIDERATIONS

All animal requisitions, housing, treatments, and procedures were conducted in accordance with applicable state and institutional laws, guidelines, and regulations, and in compliance with Directive 2010/63/EU on the protection of animals used for scientific purposes. The study was approved by the Ethics Committee for Animal Research at the University of Navarra and the Government of Navarra (Project code: 49/23; approval date: August 25, 2023). Authors adhered to recognized animal research reporting standards, including the ARRIVE guidelines.

## Notes

### Competing Interest Statement

The authors have declared no competing interest.

